## Supplementary figures and images for "Structural manipulations of a shelter resource reveal underlying preference functions in a shell-dwelling cichlid fish"

### Supplementary Figure 2

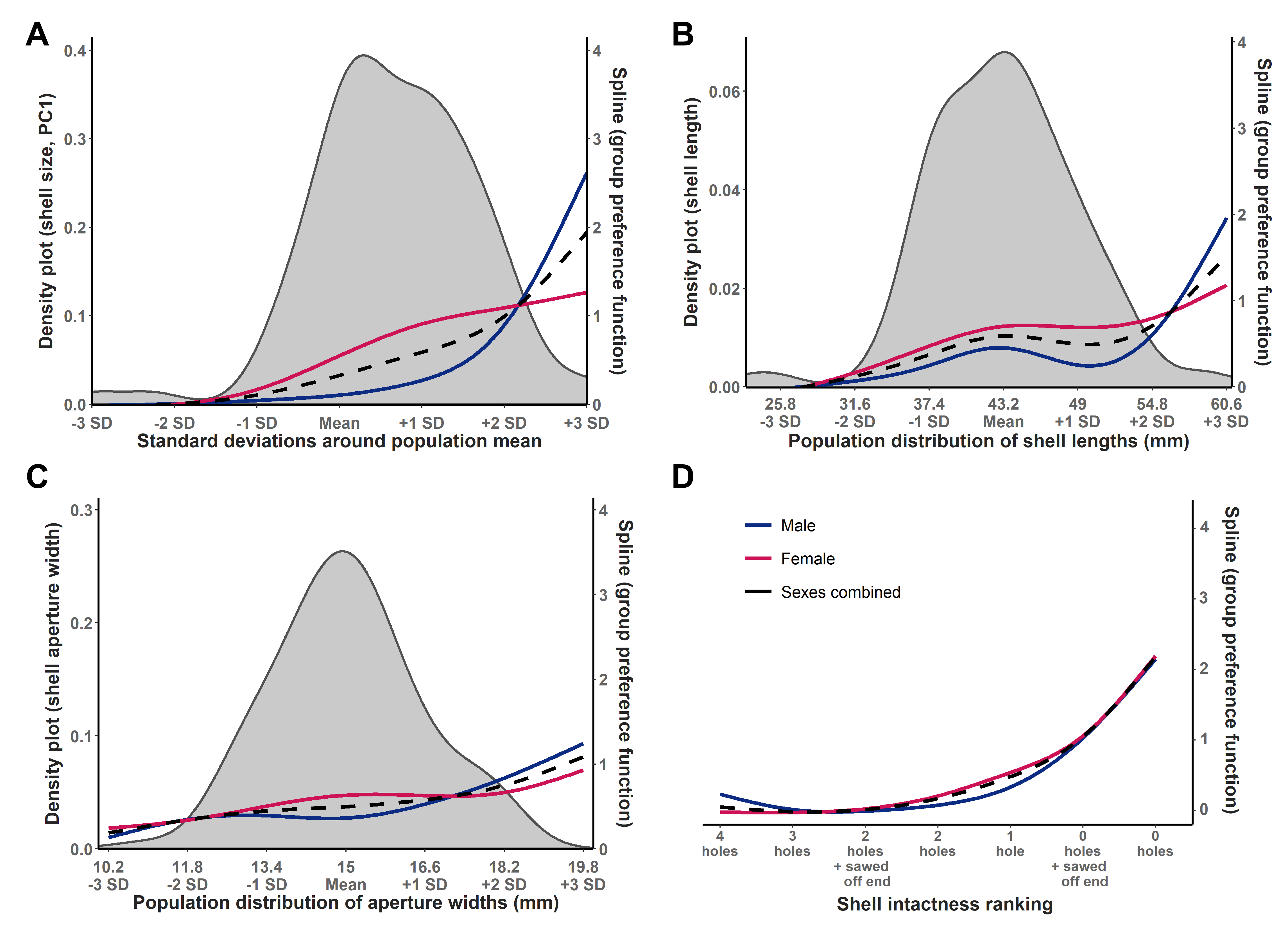

### Supplementay Figure 1

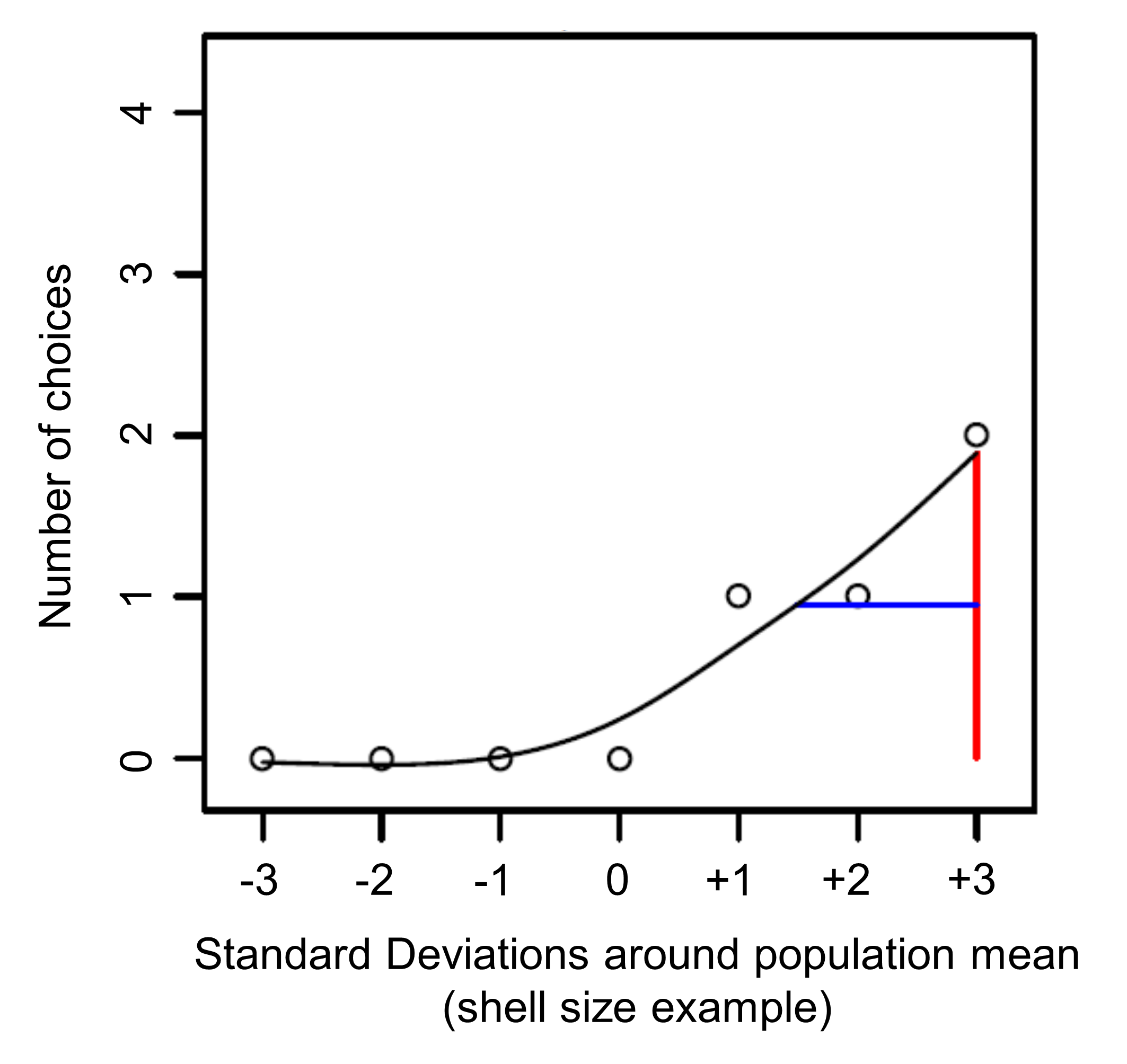
